# Spatiotemporal Dynamics of Stream Fish Assemblages Over Four Decades

**DOI:** 10.1101/592469

**Authors:** Zachery D. Zbinden

## Abstract

Fish assemblage structure was compared between 1974 and 2014 at 33 sampling locations in the Muddy Boggy River drainage, OK. The two main objectives for this comparison were to first quantify change in assemblage structure, and, second, to test for a relationship between compositional change and spatial scale. Spatial scale was manipulated by pooling assemblage data into groups ranging from K=33 pairs (i.e., local scale) to K=1 pair (i.e., global scale). Local assemblages varied in the degree of assemblage change over 4 decades (range=10-99% dissimilarity; mean=66%). The global assemblage remained quantitatively and qualitatively similar: most species persisted through time and those that did not were rare, and several diversity measures were not significantly different between time periods. Pooling assemblage data into consecutively larger groups and assessing the resultant compositional change revealed interesting patterns indicative of domains of spatial scaling. A discontinuity in the relationship between compositional change and spatial scale occurred at K=15, and this grouping is roughly the size of the headwater/tributary drainages of the Muddy Boggy system. This result suggests assemblages are more predictable when defined at scales larger than a stream reach, and a domain of higher predictability exists at a scale smaller than the entire drainage.

## INDRODUCTION

The study of communities—and assemblages within them, e.g. fish or macroinvertebrates (Fauth et al., 1996)—has largely been driven by dynamics of their composition. It has evolved considerably from its initial focus on equilibrium, as mediated by interspecific competition (Elton, 1946) and resource partitioning (Hutchinson, 1957; Hardin, 1960). Here, the driver was ecological succession, with its eventual maturation of “pioneer communities” (Shelford, 1911; Odum and Odum, 1959). With a focus on extrinsic forces and random perturbations, a distinction was made between resilience (“persistence of relationships”) and stability (how fast the system returns to equilibrium) (Holling, 1973). In this way, communities could maintain persistent groups of species despite fluctuations in abundance. Further interest in environmental perturbation envisioned competition differently and led to more complex explanations of equilibrium and species coexistence (Wiens, 1977), although these ideas were conceived earlier (Hutchinson, 1953). Following this were many investigations of the role disturbance plays in community dynamics (Harrell, 1978; Ross, 1983; Townsend, 1989). While some studies of fish assemblages demonstrated evidence for persistence (Smith and Powell, 1971; Matthews, 1986), studies from other systems called this in to question (Herbold, 1984; Yant, 1984), which provoked debate about the role of stochastic and deterministic forces shaping community dynamics and whether or not equilibrium states were really prevalent (Grossman et al., 1982, 1985; Rahel et al., 1984). Another more centrist approach emerged with the acknowledgement that indeed community structures changed over time, yet with the existence of a stochastic (May, 1973) or loose equilibrium (DeAngelis et al., 1985) as an overarching central tendency.

Around the time some ecologists were focused on pattern and the role of stochastic and deterministic processes, others were attempting to integrate perspectives and focus more broadly on landscapes, heterogeneity, and scale (Risser et al., 1984; Meentemeyer, 1989). Aside from differences inherent to various systems/drainages responsible for conflicting patterns (i.e. persistent or not), the hierarchical nature of persistence pointed to scaling (e.g. temporal, spatial, and analytical) as a potentially overlooked, confounding factor that might resolve the debate, or at least shift the focus (Frost et al., 1988; Turner et al., 1989a; Rahel, 1990). Fields outside ecology were already recognizing the importance of sampling unit size on variation of measurements (Robinson, 1950), and that variation was usually inversely related to the size of the grain/sampling unit (Meentemeyer, 1989). Previous simulation studies had demonstrated a positive relationship between spatial heterogeneity/scale and population persistence (Reddingius and den Boer, 1970; Roff, 1974; Paine and Levin, 1981). These studies, in part, inspired subsequent focus on “landscape ecology” (Galzin, 1987; Sherry and Holmes, 1988; Turner et al., 1989b) and metapopulations (Taylor, 1988; Hanski, 1994)—eventually leading to the metacommunity concept (Leibold et al., 2004). The main conclusion of this body of work was that instability at local levels resulting in extirpations need not translate to instability at broader scales when sampling units are connected by dispersal. Furthermore, fine-scale instability may even be requisite for stability at broader scales (DeAngelis and Waterhouse, 1987).

Concomitant with greater emphasis on scaling was a call to develop nonarbitrary methods for defining scales. The necessity for this was exemplified by the arbitrary choice of spatial scale in many studies, which was based more on convention than science (Frost et al., 1988). One approach, for example, was to determine the minimum area in which a community is stable and/or persistent (Connell and Sousa, 1983). Another approach was to test for “domains of scale” (Wiens, 1989) or non-linear relationships between some quantity of interest and a scaling parameter (e.g. spatial grain), which would suggest a hierarchy within which generalizations could be made regarding causal factors at different levels (O’Neill et al., 1986). Discontinuities in a relationship between a parameter and scale, and/or peaks of unusually high variance, could indicate where domains arise on the scaling axis (Greig-Smith, 1979).

A rich literature exists on spatial dynamics of aquatic communities relating to longitudinal gradients (Vannote et al., 1980; Schlosser, 1987). However, relatively few studies have evaluated the influence of spatial variation on temporal dynamics of stream fish assemblages. Those that have did not deal with spatial scale per se, but rather longitudinal gradients (i.e. stream hierarchy) (Schlosser, 1982) or differences among habitats (Bart, 1989). The studies that have explicitly considered spatial scale demonstrated greater persistence of fish communities at the broadest, compared to the finest, spatial scale (Ross et al., 1985; Matthews and Marsh-Matthews, 2016). Recent syntheses of long-term data of compositional change (Matthews and Marsh-Matthews, 2016, 2017) have compared patterns at two scales: “local” and “global” (Collins, 2000). In these cases, local was defined as the dynamics of the sampling unit/stream reach, and global was defined as the total assemblage of all local sites pooled together.

These considerations apply directly to the two main objectives of this study: I. To quantify over four decades the stability and persistence of fish assemblages in an Oklahoma river drainage, and II. To quantify how spatial scale impacts the degree of assemblage change through time. This will provide an empirical test of the “domains of scale” hypothesis (*sensu* Wiens, 1989) by defining the relationship between compositional change and spatial scale. In this sense, interpreting the way scale and change are associated will be compounded if their relationship is linear (i.e. continuous), whereas discontinuities will support domains of scale that emphasize different causal factors acting at different spatial scales. In addition, if domains do indeed exist, their spatial scale may serve to objectively define assemblage and to determine the minimum scale necessary to obtain maximum predictability.

## METHODS

The Muddy Boggy River in the Cross Timbers Ecoregion of southeastern Oklahoma, U.S.A. is a major tributary to the Red River (Pigg, 1977) draining 6,291 km^2^ from an area 113 km (north-south) by 48 km (east-west). There is little urban development and only modest manufacturing. The three largest cities in the area had the following populations as of the 2010 U.S. census: Ada (16,810), Atoka (3,107), and Coalgate (1,967).

The river flows southeastward and is formed by the confluence of Clear Boggy Creek (west – HUC 11140104) and Muddy Boggy Creek (east – HUC 11140103). Geologically, the upper reaches are in the northern Arkoma basin between the Arbuckle mountains (west) and the Ouachita mountains (east). This upper section is more rugged, and streams are higher gradient than those in the Dissected Coastal Plains near the confluence with the Red River (Pigg, 1977). The higher gradient streams are generally clear, swift-flowing, and contain riffle-run-pool structure with gravel substrate (Pigg, 1977; pers. obs.). Lower gradient streams to the south are more turbid and sluggish, and often have more homogenous, mud-bottom-channel habitat littered with coarse woody debris (Pigg, 1977; pers. obs.).

Beginning in 1974, Jimmie Pigg made 277 fish collections in the drainage, and of these, 174 were made with seine nets and in streams, while the rest used gill-nets or electroshocking equipment and/or were made in ponds or ditches (Pigg, 1977). In 2014, I made 65 fish collections in streams of the drainage (Zbinden and Matthews, 2017) during the same months (May – September) using the same gear (4.57 m × 1.22 m × 4.88 mm mesh seine) and methods (sampling all micro-habitats with 100 m stream reach) as Pigg. Efforts were made to revisit the exact stream reaches sampled four decades earlier, but changes in land use and access were problematic. All sites (174 + 65) were mapped using latitude and longitude, and each location sampled in 2014 was paired with the nearest site from 1974 (by fluvial distance). Pairs were retained for analysis if the straight-line distance between was less than eight km and the stream order was the same for both sites. Similar approaches of comparing assemblages through time have been used previously (Matthews and Marsh-Matthews, 2015). Eight of the sites sampled in 2014 were sampled again in 2015 during the same months to provide context for assemblage fluctuation over one year.

All statistical analyses were performed with R version 3.4.1 (R Development Core Team, 2016). Prior to analyses, *Campostoma anomalum* and *Campostoma spadiceum* collected in 2014 were collapsed into the group *Campostoma spp*. because recognition of *C. spadiceum* did not occur until 2010 (Cashner et al., 2010) thus were not differentiated by J. Pigg. In the same manner, *Fundulus notatus* and *Fundulus olivaceus* were collapsed into *Fundulus spp*. due to identification issues in this region (J.F. Schaefer and W.J. Matthews, pers. comm.). Finally, individuals identified as *Notropis rubellus* in 1974 were re-assigned to *Notropis suttkusi*, following Humphries and Cashner (1994).

Fish species abundance data from 1974 and 2014 were compiled into a single matrix, and the dissimilarity among all pair-wise combinations was quantified using Morisita-Horn index (Morisita, 1959; Horn, 1966; Jost et al., 2011; Matthews and Marsh-Matthews, 2017) with the R package ‘vegan’ (Oksanen, 2015). Only distances between target pairs (i.e. the matching local sites from ’74 and ‘14) were extracted from the dissimilarity matrix. Note, Morisita-Horn index is typically presented as a ‘similarity’ index, meaning it ranges from 0 to 1, or no overlap to complete assemblage overlap respectively. However, the methods available within the function ‘vegdist’ of the R package ‘vegan’ are adjusted to be dissimilarities, meaning the opposite is true (i.e. 0 = complete overlap and 1 = total dissimilarity); in other words, the diagonal values across the matrix between each assemblage and itself will always be equal to 0. Therefore, I will refer to “dissimilarity” and “Morisita-Horn dissimilarity” throughout.

Non-metric multidimensional scaling (NMDS, Kruskal, 1964) was used to visualize the dissimilarity matrix in multivariate space. This visualization allows inspecting differences among local assemblage pairs and the differences among global “clouds” of assemblages from 1974 and 2014. In addition, NMDS containing two sets of samples from different time periods allows for visualization of parallel trajectories among sites within the drainage. Stress less than 0.20 was considered the threshold for accurate representation of the data (Kruskal, 1964). To test for the effect of geographic distance between assemblages and time between collections, dissimilarity values were pooled into three groups: 1974-2014 pair-wise comparisons from the precisely same location (n=12); 1974-2014 pair-wise comparisons between approximately matching locations (n=21); and pair-wise comparisons between 2014 and 2015 (n=8, all exact matches). Linear regression was used to test the relationship between Morisita-Horn dissimilarity and geographic distance between appropriate site pairs (both straight-line and fluvial distance).

Diversity of the global assemblage (33 sites pooled) was compared between 1974 and 2014 using a variety of measures. The R package ‘diverse’ (Guevara et al., 2016) was used to calculate mean species richness (MacArthur, 1965) and Simpson’s reciprocal diversity (Simpson, 1949). Differences between sampling periods were analyzed using Student’s *t*-test. The R package ‘vegetarian’ (Jost, 2007; Charney and Record, 2015) was used to compute alpha, beta, and gamma diversities. Values were obtained via bootstrapping with 10,000 iterations.

At the finest spatial level, or the “local” level, the data set contains 33 pairs of sampling localities where fish were collected in 1974 and 2014. At the broadest spatial level, or the “global” level, the data contain 1 pair of pooled localities sampled in 1974 and 2014. The R package ‘ClustGeo’ (Chavent et al., 2018) was implemented to create a hierarchy based on spatial location to create the intermediate groupings between 33 pairs and 1 pair. A tree of all sampling localities was created using function ‘hclustgeo’ which implements a Ward-like clustering algorithm. The algorithm requires two distance matrices as input, in this case: Morisita-Horn dissimilarity among sites and Euclidean distance between sites. However, alpha weight (1.0) was used as the mixing parameter so that the spatial matrix alone would be used to cluster the sites. Thus, all 33 sites were clustered based on spatial proximity to one another. The function ‘rect.hclust’ was used to visually identify clusters of sites on the dendrogram for each cluster value K from K=33 to K=1. A separate community matrix was created for each clustered group containing the pooled data of the sites within each group for 1974 and 2014. For example, for K=2 there were 4 rows of species abundance data: 2 sites X 2 sampling periods. Just as described above, Morisita-Horn dissimilarity index matrices were calculated for each of the 33 community matrices. The compositional distances between 1974 and 2014 for each group within a clustered set was then extracted. So, for each group K there would be K number of distances. The distributions of the distances for each spatial cluster were visualized using R package ggplot2 (Wickham, 2010) to visually inspect the relationship between the number site pools (i.e. spatial scale) and the compositional change over 40 years (M-H index).

## RESULTS

Fish assemblages from 33 locations (Fig. 1) sampled in 1974 and 2014 were compared. Local assemblages showed considerable variation in the degree of compositional change between sampling periods, but overall composition at the global level was little changed. There were 37 species collected during both 1974 and 2014, 9 species were collected only in 1974, and 7 species collected only in 2014 (Appendix 1). All species not collected during both sampling periods were rare: occurring at only one site (n=14) or at an abundance of one individual per site (n=2). None of the global diversity indices differed between sampling periods: Simpson’s Reciprocal Diversity (1974= 4.23 ± 0.80 SE and 2014= 3.95 ± 0.32), Alpha diversity (1974= 9.48 ± 0.15 and 2014= 10.79 ± 0.14), Beta diversity (1974= 4.85 ± 0.20 and 2014= 4.17 ± 0.11), Gamma diversity (1974= 46.0 ± 1.60 and 2014= 45.0 ± 0.86).

**Figure 1.**
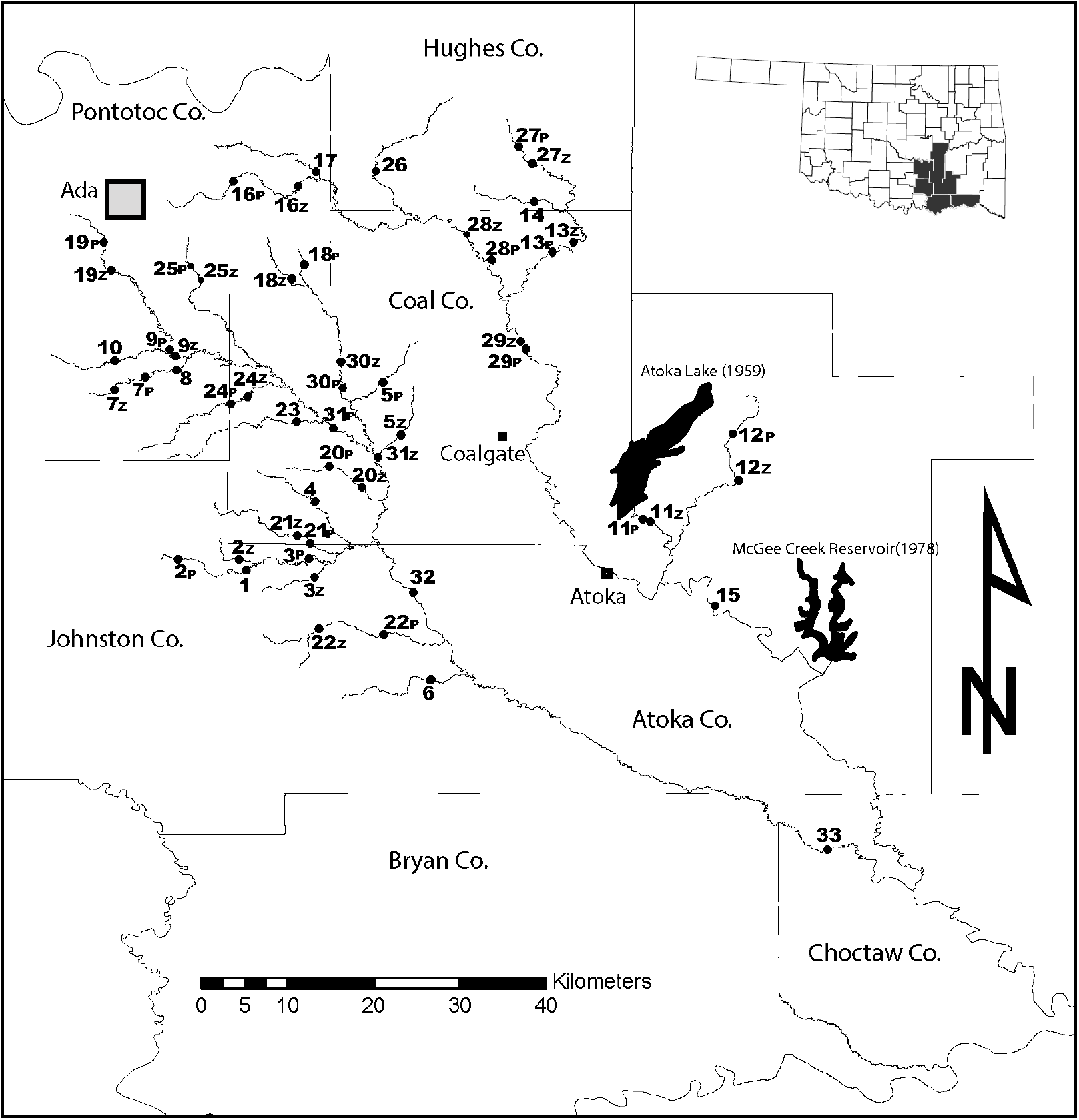
Map of sampling locations in the Muddy Boggy River drainage. If the locations were not identical reaches, each is denoted by a point and number followed by either a “P” for “Pigg 1974” or a “Z” for “Zbinden 2014”.

Local assemblages often changed drastically between 1974 and 2014. Local assemblages ranged from 10% to 99% dissimilarity, and mean dissimilarity was 66% for the 33 pairs. Assemblages changed more between 1974 and 2014 than they did between 2014 and 2015 (mean = 66% and 35%, respectively; t-stat = 3.32, n = 8, p = 0.002). Assemblage dissimilarity did not differ between groups of exact location/same stream reach (mean= 66%) and approximate location matches (mean= 66%), nor was dissimilarity for matching sites correlated with straight line (*R^2^* = 0.030, p = 0.339), or fluvial distance between sites (*R^2^* = 0.028, p = 0.354). This suggested the scheme used to select sites for comparison had not influenced the analyses. Fig. 2 illustrates the distributions of dissimilarity values of the 40-year comparison, 1-year comparison, exact matching site comparison, and approximate site comparison.

**Figure 2.**
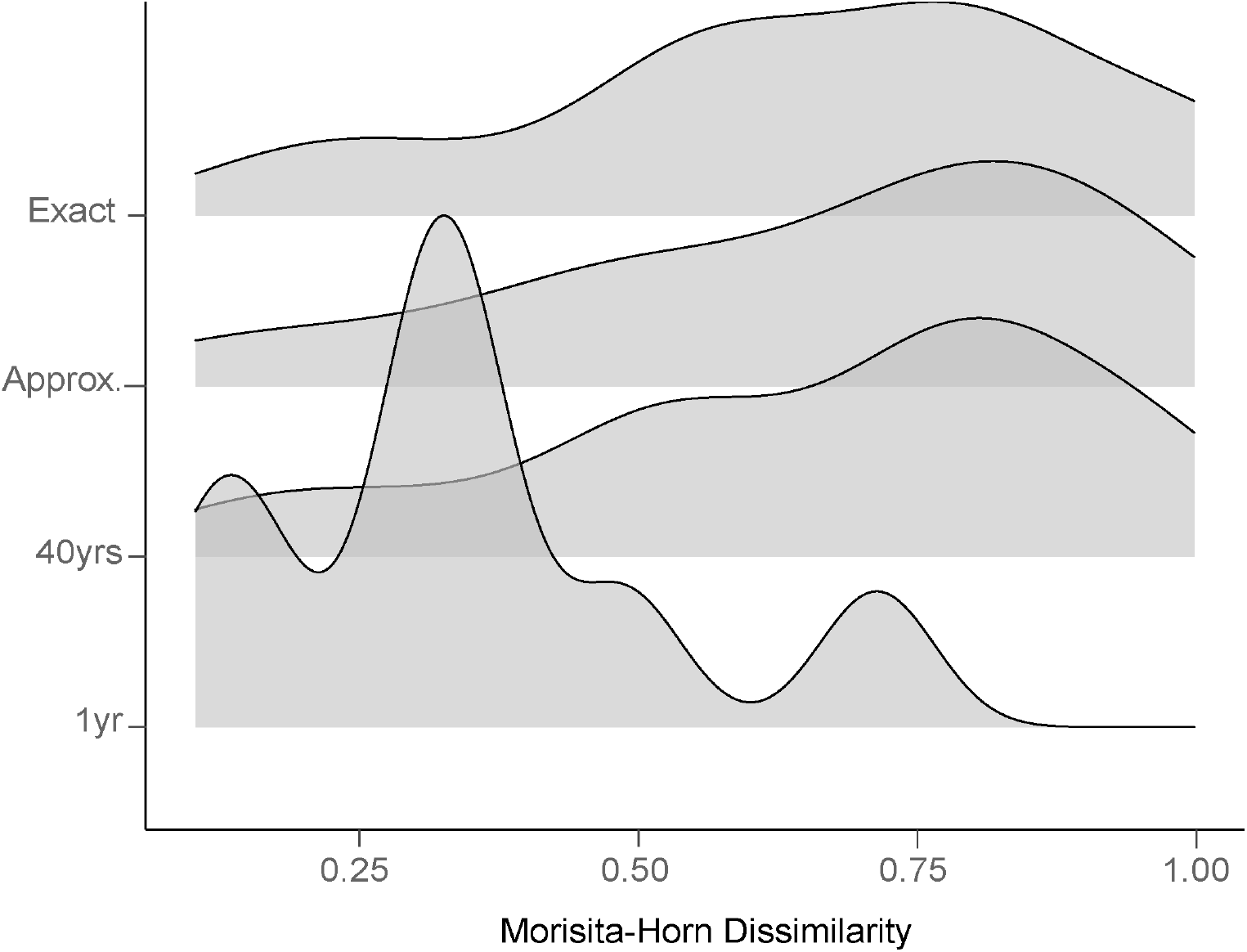
This figure illustrates the distribution of compositional change among groupings of interest including: I) 40-year comparisons of sites that were exact geographic matches, II) 40-year comparisons of sites that were approximate geographic matches (distance between ≤ 8 km), all 40-year comparisons together, and 1-year comparisons between 2014-2015.

NMDS produced 3 axes using 100 runs with a stress = 0.175. Fig. 3 provides an illustration of assemblage variation through time via NMDS, which highlights the findings from the diversity analysis presented above. The “clouds” of points from 1974 and 2014 were similar in location and spread. Panels B and C of Fig. 3 demonstrate two points: first, assemblages varied in how much change occurred through time no matter if the sites were the same reach (Panel C) or approximate matches (Panel B), and second, there is no pattern of parallel trajectories that would suggest common shifts in composition across sites.

**Figure 3.**
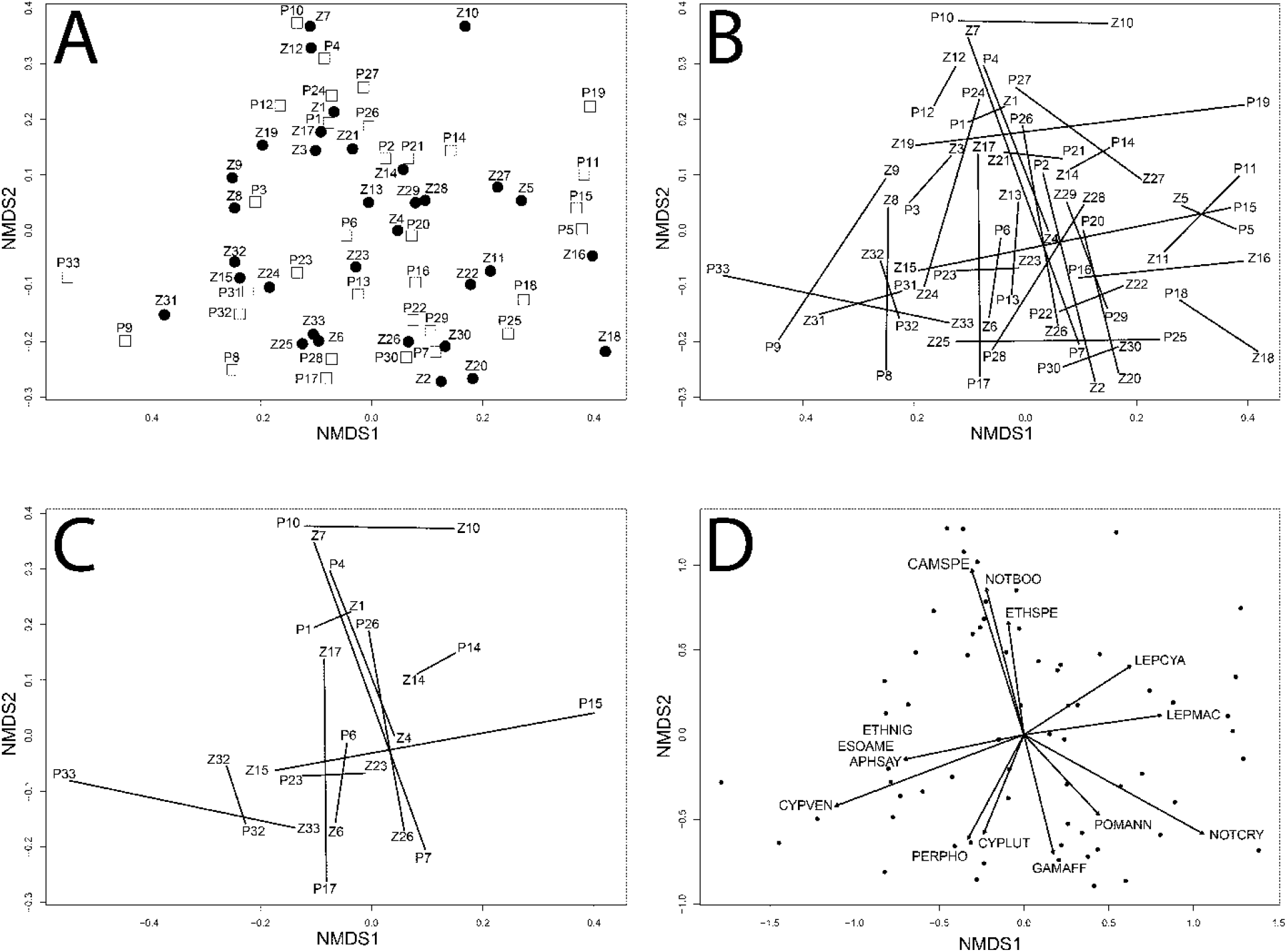
A) NMDS of 33 assemblages from 1974 (white squares labelled “P”) and 33 assemblages from 2014 (black circles labelled “Z”). **B)** The same NMDS from A, but with vectors illustrating assemblage change among sampling periods; **C)** NMDS with only the 12 exact geographic matching sites; **D)** Ordination of NMDS with species significantly related to axes shown as vectors. Species identities are coded as the first three letters of the genus and species name. See Appendix 1 for additional species codes.

The relationship between number of site clusters and the compositional change over 40 years is shown as a boxplot in Fig. 4. Note, when sites are clustered into one group (K=1) there is no variation and so only one distance value for the comparison between years is shown (i.e. the “bar” above K=1). At the global level (K=1), which includes all sites sampled, the MH-dissimilarity between the two sampling periods was 19% and the mean MH-dissimilarity for the comparison at the finest-scale (33 pairs) was 66% dissimilarity. The distribution of dissimilarity values decreases in median value as the clusters get larger, or as more sites are pooled together (moving right to left on the x-axis of Fig. 4). A sharp decline in the median distribution begins at approximately k=20, before reaching a local minimum at k=15. For an idea of the spatial scale of pooled sites at this local minimum see k=15 panel of Fig. 5. Following the decrease, the median begins to vary widely as the clusters grow larger in size. In addition, the variance of the distribution, illustrated by the interquartile range, also appears to change with cluster size.

**Figure 4.**
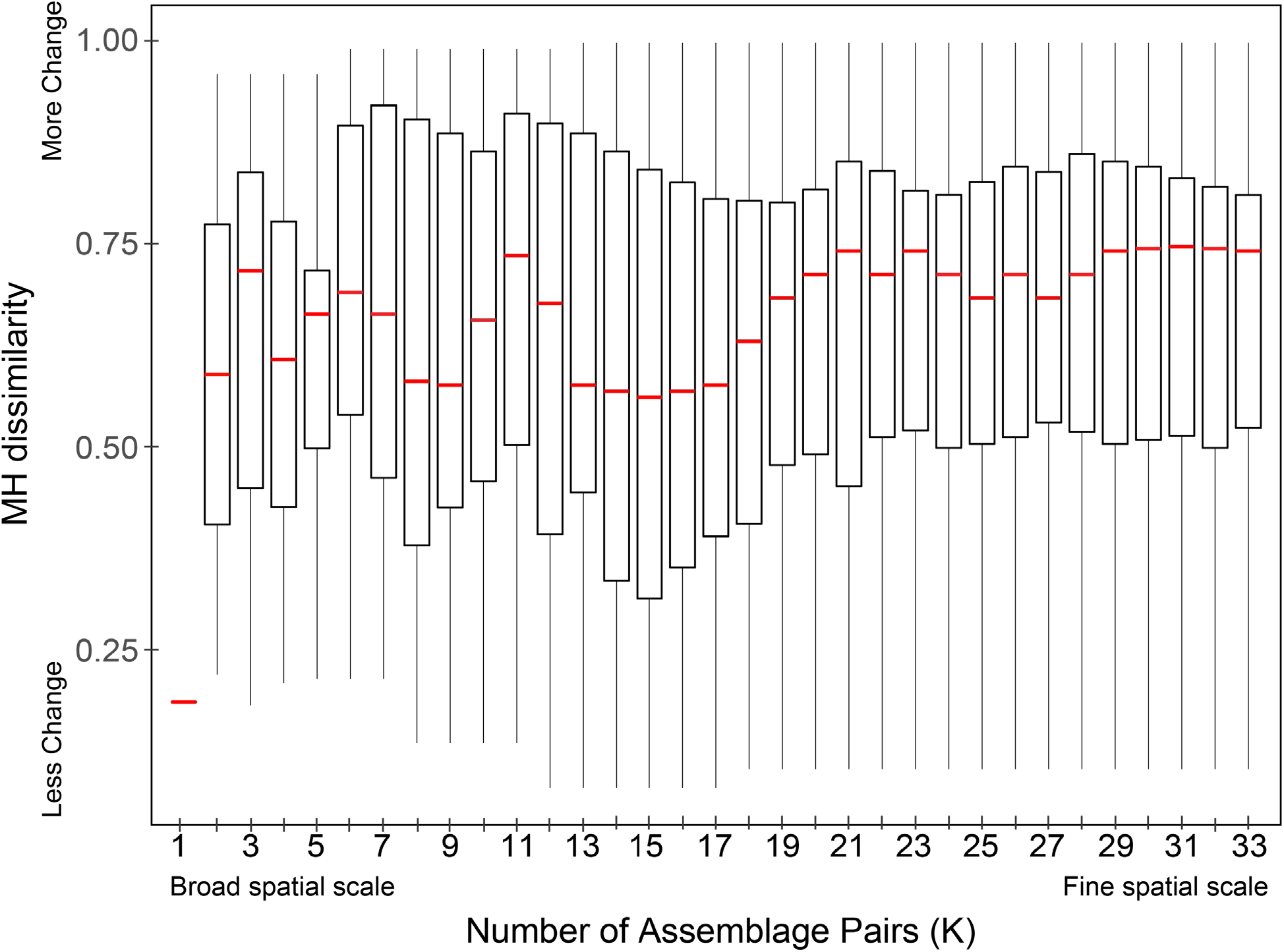
A boxplot showing the relationship between the number of the spatial clusters (K) and assemblage compositional change between 1974 and 2014 (Morisita-Horn dissimilarity) for the sites within the clusters. Red lines indicate median dissimilarity values.

**Figure 5.**
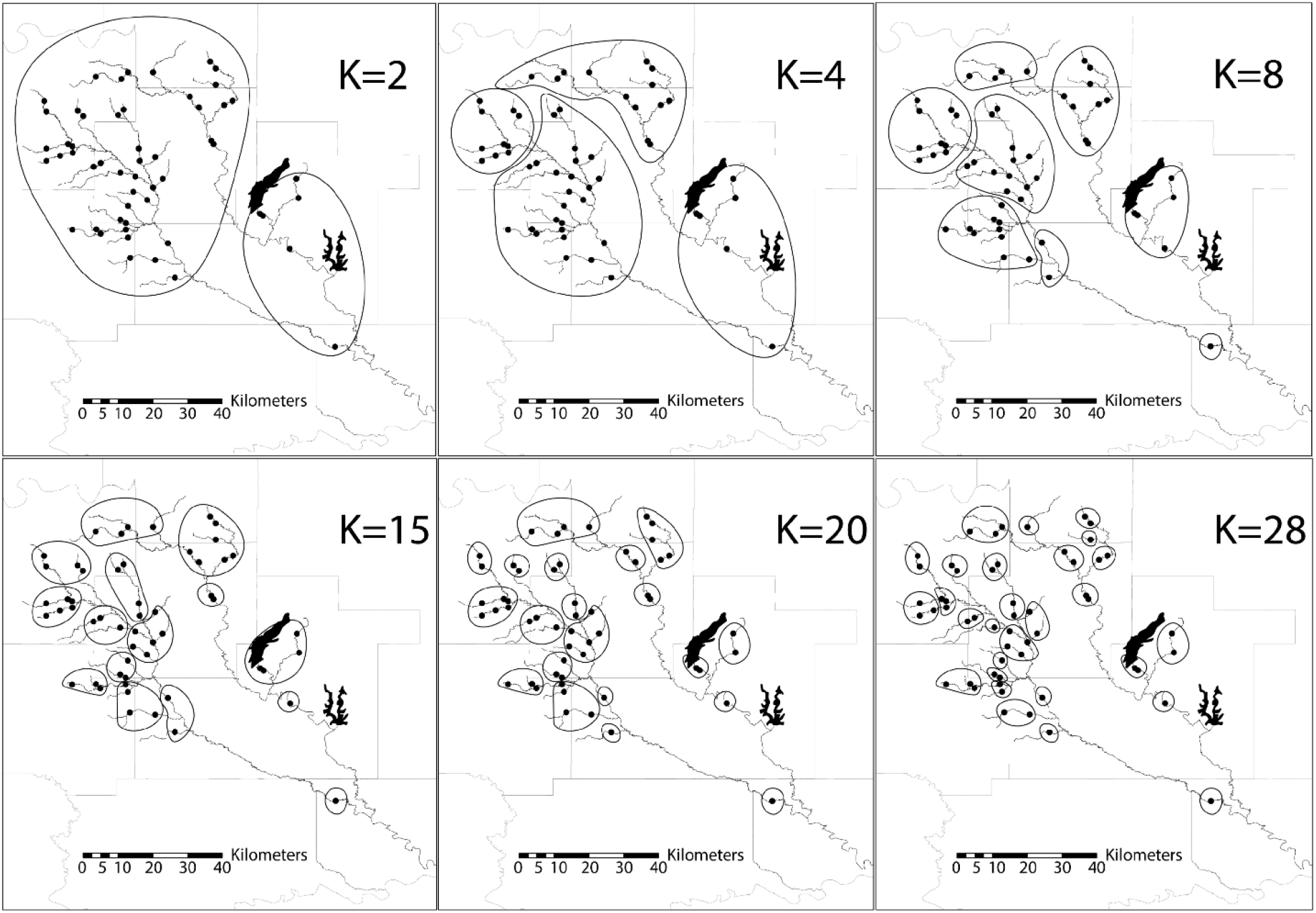
These six maps of the drainage show examples of spatial clusters of sites determined using the clustering algorithm. For the analysis all spatial clusters from K=1 to K=33 were created.

## DISCUSSION

This study highlights the importance of spatial scale regarding temporal change in assemblages. The Muddy Boggy fish assemblage remained qualitatively and quantitatively similar across 40 years at the global/drainage scale. There were 37 species present at both sampling periods, and those present in only one sampling event were rare. Many common species had striking similarities in total abundance across 40 years (Appendix 1), for example: *Notropis boops* 719 – 668; *Cyprinella lutrensis* 654 – 640; *Cyprinella venusta* 573 – 546; *Pimphales notatus* 132 – 144. There were large declines in abundance for some species (e.g. *Etheostoma radiosum, Notropis nocturnus, Ictalurus punctatus*, and *Lepomis cyanellus*), while there were increases in others (e.g. *Campostoma spp., Pimephales promelas, Notropis suttkusi, Lepomis humilis*). Measures of mean alpha diversity corroborate the persistence of diversity over time, and a lack of change in beta diversity measures suggest this system has not experienced increased homogenization of fish taxa since the 1970s. This lends support to the conclusion of Ross (2013) that most systems tend to have high levels of persistence and stability over time. In addition, these results agree with the synthesis of Matthews and Marsh-Matthews (2017) which included 40+ years of repeated sampling of drainages in Oklahoma and Arkansas (Matthews et al., 2013; Matthews and Marsh-Matthews, 2016). The long-term data presented by those authors suggested high persistence of species assemblages, and when instability occurred from year to year, assemblages tended to return to the original structure. They concluded the Loose Equilibrium model was the best explanation for assemblage change in their study systems. The dataset presented here only contained samples at two time periods, which does not allow for analysis of trajectories (Matthews et al. 2013), but based on the persistence at the global level over four decades, the results of this study support the notion of a loose equilibrium.

Local assemblages, however, varied considerably between 1974 and 2014. Mean percent dissimilarity was 66% at the local scale (n=33 site pairs) and some sites were as much as 99% dissimilar, which indicates complete turnover in species composition. This result mirrors that of other studies that found variability at the local scale while measuring a lack of change at the global scale (Ross et al., 1985; Matthews, 1986; Magurran and Henderson, 2010; Matthews et al., 2013). Differences in assemblage dynamics over time at broad versus fine spatial scales may be due to the relative importance of different factors which govern the dynamics at these levels (Wiens, 1989). For example, local dynamics might result largely from colonization and extirpation due to deterministic processes, such as competition and predation, and perturbations, such as drought and floods. The dynamics at broader scales like the entire drainage may be coupled more with regional species pools and climate.

Other studies of spatiotemporal assemblage dynamics have focused on the difference in compositional change over time between headwater and downstream reaches or across stream hierarchy (Schlosser, 1982; Ross et al., 1985; Taylor and Warren, 2001; Hitt and Roberts, 2012). Generally, the prediction is that assemblages of headwater streams will tend to change more over time compared to downstream reaches due to high environmental variability of headwaters (Schlosser, 1982; Matthews, 1998), and also due to differences in characteristics and life histories of species which occur in headwaters (Hitt and Roberts, 2012). This pattern, however, may not be as general as previously thought. Analyses of long-term fish assemblage change across 9 different systems (Matthews and Marsh-Matthews, 2017) showed a longitudinal pattern in compositional change over time in 5 of the systems, but in 3 of the 5 it was the opposite pattern predicted (headwater sites showed the most stability). To test for longitudinal patterns of compositional change in the present study, linear regression was employed between Morisita-Horn dissimilarity (n=33) and several factors including stream order, maximum stream width, latitude, longitude, and percent riffle habitat. These analyses yielded no significant relationships, even without a correction for multiple tests. This is not to say spatial patterns are not occurring in the Muddy Boggy system, but perhaps these patterns themselves change and shift over time (Ross et al., 1985), and so are best measured over shorter timespans.

Fig. 4 shows three apparent declines in median Morisita-Horn dissimilarity going from fine spatial scale to broad spatial scale which are then followed by variability. The first decline occurs at K=27, the second at K=15, and the third at K=1. Much of the variability in median MH dissimilarity, especially between K=12 and K=2, may be due to high dissimilarity found in the sites in the southeast corner of the drainage, below two reservoirs (Sites 11, 12, 15, and 33). Site 33 in particular was 99% dissimilar between sampling periods, and when all of these sites were included in the same pool (K=5) the group was 96% dissimilar between sampling periods. Although Atoka lake was constructed 15 years before Pigg’s collections, McGee Creek Reservoir was constructed just 4 years after. These data lack the power to determine if the reservoirs affected assemblage dynamics, but this question could inspire future study designs of long-term assemblage investigations.

The sharpest decrease in assemblage change as it relates to spatial scale occurs at K=15. The size of the groups at this cluster size (Fig. 5) is roughly the size of the headwater drainages (Stream Order ≤ 3) which are tributaries to the main channel of either Clear Boggy or Muddy Boggy Creek. This relationship supports the “spatial domains” hypothesis regarding zones along the spatial continuum that occur because of a shift in the importance of the extrinsic and intrinsic forces that shape assemblage dynamics between zones (Wiens, 1989). The relationship between scale and temporal dynamics appears to contain discontinuities, and when discontinuities appear, they are followed by unusually high variation (Greig-Smith, 1979; O’Neill et al., 1986).

Another study of fish assemblage change over time (Hoeinghaus et al., 2003) found discriminant function analysis could assign the identity of a creek (i.e. tributary) based on the fish assemblage found there, which suggested creek-specific fish assemblages. These creek-specific assemblages may become isolated by the main channel of the larger drainage where there are obvious changes in habitat, depth, and predator densities which may limit dispersal between creeks. Therefore, the isolated tributary systems of larger order streams such as Clear and Muddy Boggy (4^th^ to 5^th^ order) may be modelled as meta-communities (Leibold et al., 2004; Muneepeerakul et al., 2008) influenced by the characteristics intrinsic to the sub-drainage. At this spatial scale dynamics become more predictable because the system is partly closed to immigration and emigration, which in part makes prediction at the scale of an individual stream reach so difficult. Most species of fish in southeastern Oklahoma readily move in and out of a 100m stream reach over a lifetime, but moving out of a tributary drainage is more difficult (Radinger and Wolter, 2014). Systems such as Brier Creek, OK and Piney Creek, AR, noted above (Matthews et al., 2013; Matthews and Marsh-Matthews, 2016), are much smaller than the Muddy Boggy system but compare closely to the size of the groupings shown at K=15 (i.e. tributary drainages). This evidence suggests assemblages are more predictable when defined as a group of fish occurring within an area much larger than a single stream reach, possibly the size of a 2^nd^ or 3^rd^ order tributary drainage.

The fish diversity found across the Muddy Boggy River drainage has been persistent since Jimmie Pigg surveyed the region 40 years ago. Fluctuations have occurred at any given stream reach, and in some cases an entirely different group of fish may now be found in some reaches. Assuming the meta-community model is good at explaining stream fish assemblages, then perhaps Connell and Sousa (1983) were correct that instability at local scales is essential for the stability of the larger system. As predators, competitors, or drought cause declines in populations once inhabiting a sampling station, there may indeed be opportunity for new populations to arise elsewhere in the environmentally heterogeneous system. The spatial scale at which these dynamics become relatively stable and therefore predictable over timescales relevant for management remains an important question. Hopefully more studies of long-term community change will explore these patterns related to spatial scale.

## ACKNOWLEDGMENTS

Financial support for this study was provided by the Oklahoma Department of Wildlife Conservation through the State Wildlife Grant FI 3AF01213 (T-74-1) and the University of Oklahoma. Fish sampling was done under the approval of the IACUC of the University of Oklahoma (R14-002) and with scientific collecting permits from Oklahoma Department of Wildlife Conservation (No. 5995 & No. 6166). I thank W. Matthews and E. Marsh-Matthews, who were primary investigators for the Oklahoma State Wildlife Grant that funded the collecting efforts for this study and helped improve this manuscript. I thank K. Montgomery and M.E. Douglas for comments and critiques. I thank A. Geheber and R. Lehrter for field assistance. Finally, I thank the late J. Pigg for his extensive field collections and contribution to the study of ichthyology in Oklahoma.

**Appendix 1.**
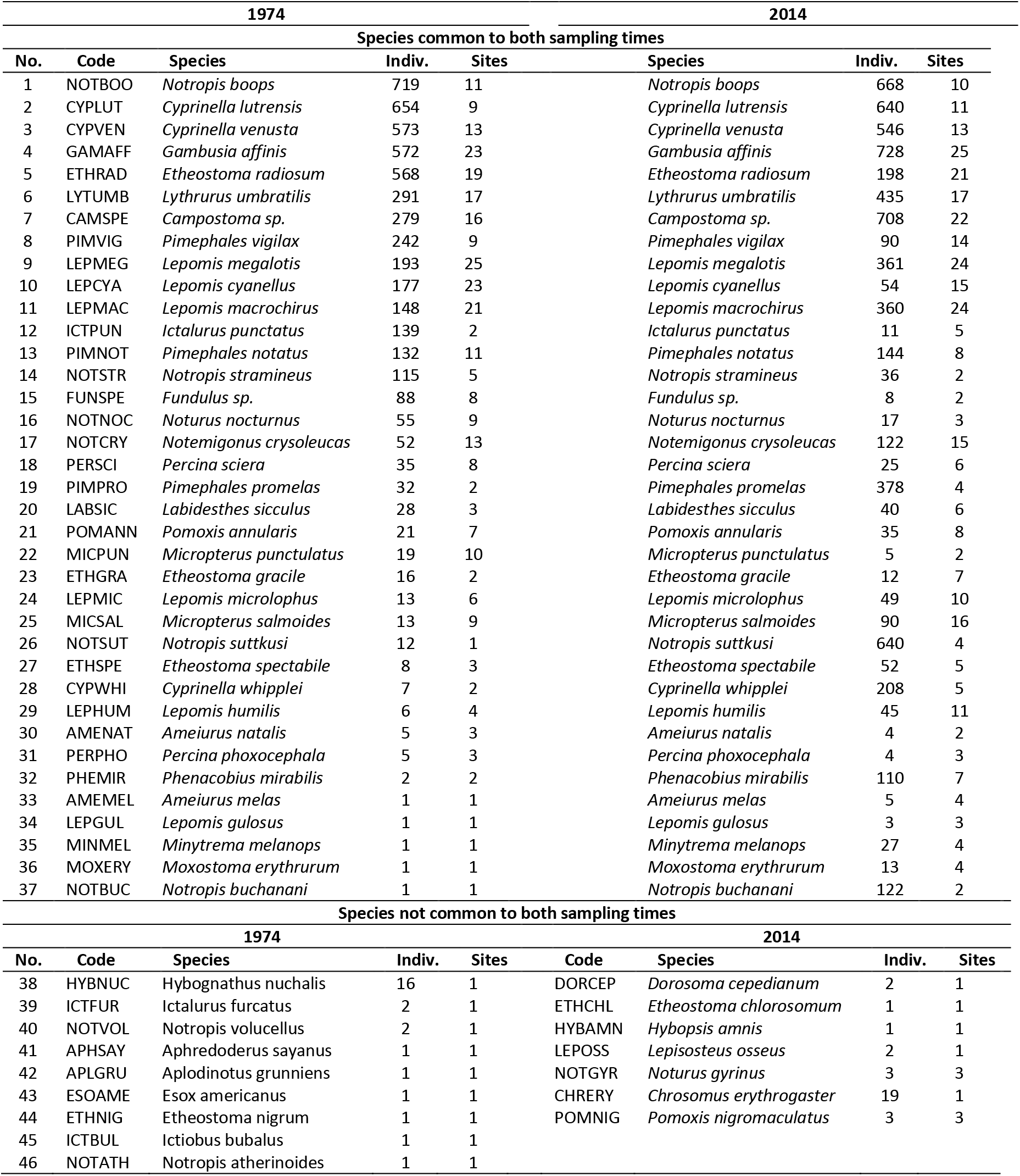
Species list for 33 sites sampled in 1974 (left) and 2014 (right). Species sorted by abundance in 1974.

